# Direct and delayed synergistic effects of marine heatwaves, metals and food limitation on tropical reef-associated fish larvae

**DOI:** 10.1101/2022.02.23.481600

**Authors:** Minh-Hoang Le, Khuong V. Dinh, Xuan Thi Vo, Hung Quoc Pham

**Affiliations:** Cam Ranh Centre for Tropical Marine Research and Aquaculture, Institute of Aquaculture, Nha Trang University, No 2 Nguyen Dinh Chieu Street, Nha Trang City, Vietnam

**Author notes:** Co-first authors (Minh-Hoang Le, Khuong V. Dinh). Corresponding author: Khuong V. Dinh.

**Keywords:** food availability, marine heatwave, pollution, tropical fish, tropical marine ecosystems

## Abstract

Tropical fish are fast-growing and high energetic-demand organisms, which can be highly vulnerable to long-lasting effects of heat stress and pollution, particularly under food shortages. We tested this by assessing highly complex direct and delayed interactive effects of an extreme temperature (32°C) from a simulated marine heatwave (MHW), copper (Cu, 0, 100, 150 and 175 µg L^-1^) and food availability (limited and saturated food) on larvae of a tropical, reef-associated seaperch (*Psammoperca waigiensis*). Cu, MHW, and food limitation independently reduced survival and growth, partly explained by reduced feeding. The negative effect of Cu on fish survival was more substantial under MHW, particularly under limited food. Delayed interactive effects of Cu, MHW, and food limitation were still lethal to fish larvae during the post-exposure period. These results indicate that reef-associated fish larvae are highly vulnerable to these dominant stressors, impairing their ecological function as predators in the coral reefs.

**Graphical abstract:** 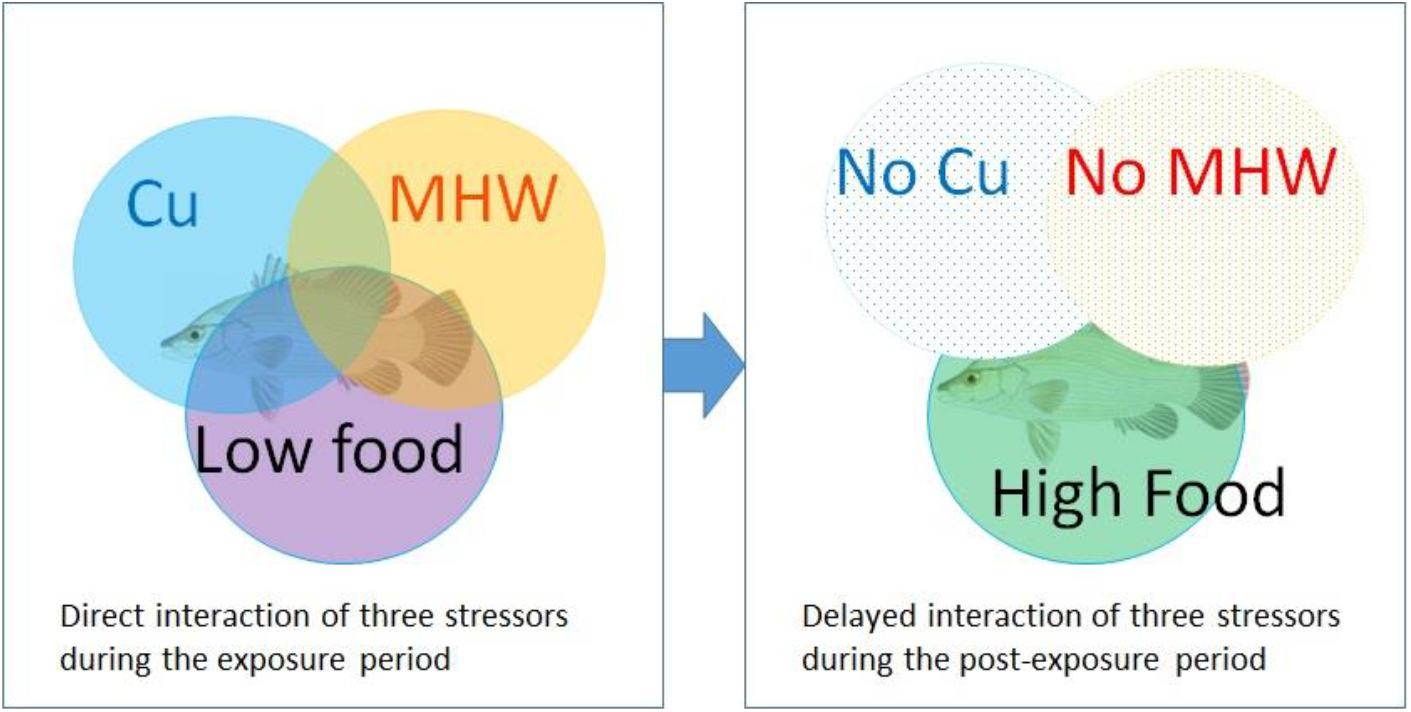

## 1. Introduction

Marine heatwaves emerge as one of the dominant threats for marine organisms (Oliver et al., 2018; Smale et al., 2019). Tropical organisms are particularly vulnerable to marine heatwaves because MHW temperatures can exceed their upper thermal optimum (Nguyen et al., 2011; Rosa et al., 2014; Tewksbury et al., 2008), resulting in rapidly reduced growth (Le et al., 2020; 2021) or even causing mass mortality (e.g., Garrabou et al., 2009), reshaping structure, and functions of typical tropical coastal ecosystems such as seagrasses and coral reefs (Dalton et al., 2020; Short et al., 2015). In tropical countries around the Indo-Pacific, millions of people are heavily based on fisheries and aquaculture for their livelihood, effects of MHWs on the fish stocks will directly threaten their income and food security. For example, aquaculture production of this region, particularly finfish, provides ∼ a quarter of the global fish production (SEAFDEC, 2017). Climate change, particularly heat stress, may reduce up to 40% of aquaculture species, particularly farmed finfish species in the tropical regions, in the next three decades (Oyinlola et al., 2020).

Another dominant stressor in tropical marine ecosystems is the widespread pollutants that can cause serious ecological consequences (Dinh, 2019; Jambeck et al., 2015), including a recent massive fish kill in coral reefs (Dinh, 2019). Among others, copper (Cu) compounds are one of the widespread metals in the coastal regions as these chemicals are used in large quantities in mariculture and maritime wastes released to the coastal environment (Ebenezer et al., 2014; Marcussen et al., 2014; U.S. Environmental Protection Agency, 2009; Van Sprang et al., 2007). Cu concentrations of 0 – 560 µg L^-1^ have been measured in the water nearby coral reefs (Jonathan et al., 2011; Le and Nguyen, 2018). Cu can be adsorbed quickly to suspended particles and sinks onto sediments within several hours to several days (Marcussen et al., 2014), which may impose short-term exposures of coastal marine organisms to Cu.

One crucial aspect that has often been overlooked in laboratory experiments is food availability, which may be a critically important factor determining the susceptibility of studied species to multiple stressors. Indeed, most multiple-stressor studies have been conducted in the laboratory under food saturation levels (e.g., Dinh et al., 2020b; Krause et al., 2017; Pham et al., 2020). However, stressor effects on life history and physiology may only be present or be magnified under food shortages (e.g., Bignami et al., 2016; Dinh et al., 2016a). In nature, food shortages are periodically widespread (Metcalfe and Monaghan, 2001), generally related to the seasonal variations in the abundance of prey species (Cohen et al., 2018). Food shortages may be more severe during MHWs as MHWs can impair the productivity of key prey species for marine fish such as copepods (Davis, 1985; Doan et al., 2019; Truong et al., 2020). Therefore, it is highly relevant to assess how food limitation may interact with MHWs and coastal pollutants such as Cu to affect coastal marine species.

The interaction of warming and contaminants has emerged as a central research theme in global change biology and ecotoxicology in the last decade (Crain et al., 2008; Dinh et al., 2020a; Dinh et al., 2021; Dinh et al., 2020b; Moe et al., 2013; Orr et al., 2020; Sokolova and Lannig, 2008). However, only a few studies have investigated whether an additional stressor, e.g., food limitation, competition, or predation stress, may further modify the interaction of warming and contaminants (Crain et al., 2008; Dinh et al., 2016a; Knillmann et al., 2013; Pham et al., 2020). For example, food limitation combined with the heatwave makes pesticides lethal to aquatic insects (Dinh et al., 2016a). Three-stressor experiments have rarely been conducted on marine species (reviewed in Crain et al., 2008).

Furthermore, multiple-stressors and ecotoxicological studies have mainly focused on the immediate effects and interactions of stressors during exposure (Crain et al., 2008; Jackson et al., 2021). However, recent studies have highlighted the role of ecological memory, delayed effects of single and multiple stressors may last several days to years post-exposure (Jackson et al., 2021). Delayed effects of stressors may affect post-exposure survival, growth, and physiology (see e.g., Dong et al., 2020; Le et al., 2020), can be carried over to the adult stage (Dinh et al., 2016b; 2016c) or from parents to offspring (Dao et al., 2018; Dinh et al., 2021; 2020b). However, we know very little about delayed effects and particularly delayed interactions of past exposures to multiple stressors on shaping the fitness of marine fish.

In this study, we tested four important research questions. 1) Whether MHW may increase the toxicity of Cu to tropical coastal fish larvae (see e.g., in tropical copepods Dinh et al., 2021; Dinh et al., 2020b), and 2) how food limitation may shape the single and combined effects of MHW and Cu (see e.g., Dinh et al., 2016a). As MHW, Cu and food limitation typically occur in short periods, we tested 3) whether and to what extent MHW, Cu and food limitation may have delayed effects; if yes, 4) how would these stressors interact to affect previously exposed fish larvae? *Psammoperca waigiensis* larvae were chosen as study animals as this species is commonly found throughout the coastal ecosystems, particularly coral reefs of the Indo-Pacific regions (https://www.fishbase.se/summary/8212) and are highly vulnerable to MHW (Le et al., 2021). *P. waigiensis* is also an important aquaculture species (Le and Pham, 2017). We quantified three major fitness-related traits for the larvae: survival, growth rate, and feeding rate.

## 2. Materials and methods

### 2.1 Ethics statement

Based on the National Regulations for the Use of Animals in Research in Vietnam: The Law of Animal Husbandry of Vietnam, 2018 and The Government Decree 32/2006/ND-CP on Management of Endangered, Precious, and rare Species of Wild Plants and Animals, experiments on waigieu seaperch are not in the list of ethical approval requirements. However, the authors have implemented their best practice following the guidelines of using animals in research based on EU directive 2010/63.

### 2.2 Experimental fish

Fish larvae were collected from a pond at Cam Ranh Centre for Tropical Marine Research and Aquaculture, Institute of Aquaculture, Nha Trang University (NTU), Vietnam. Fish were collected and acclimatized to the laboratory conditions in a similar procedure described in our previous study (Le et al., 2020).

### 2.3 Experimental design and setup

To test how MHW may affect the Cu toxicity on fish larvae and how food availability may modulate the single and combined effect of MHW and Cu, larvae were exposed to one of 2 temperatures (28 and 32°C) × 4 Cu levels (0, 100, 150, 175 µg/L) × 2 food levels (low and high) × 3 replicates. In each replicate (1-L bottle), there were 10 fish larvae, resulting in 480 individuals in total. The initial body weight and total lengths of juveniles were 160 ± 10 mg and 2.53 ± 0.11 cm, respectively. Fish was fed at a 50% saturation level (∼1.5% body weight) in the low food level. In the high food level, fish was fed at 100% saturation level (∼3% body weight).

The exposure lasted 10 days which corresponded to a critical period in the early life history of the fish and a MHW event (5 days to several weeks, Hobday et al., 2016). Survival, growth, and feeding were determined at the end of the exposure (see below for the detailed descriptions). After the exposure, larvae were transferred to the uncontaminated environment and reared for 13 days at 28°C, and all were fed *at libitum* to test to what extent they could recover from multiple stressors. Response variables were determined similar to those collected at the end of the exposure period. During the entire exposure and recovery periods, we kept salinity at 32 - 33 psu, pH 7.89 - 8.05, and dissolved oxygen > 5.0 mg/L. Fish larvae were fed with copepods and *Artermia*.

### 2.4 Data collection

Survival was checked daily. The percentage of larvae that survived to the end of the Cu exposure and the end of the experiment was determined by dividing the number of surviving larvae at each stage by the total number of larvae in each experimental unit from the start of the experiment.

Prior to the feeding trial, one larva from each experimental unit was collected and starved for 12h in the same exposure solution. Subsequently, each larva was fed with 200 adult copepods of *Pseudodiaptomus annandalei* at the end of the exposure period or 20 *Artemia* at the end of the recovery period (as fish was already quite big). The feeding trial lasted for 5 min. The fish larva was returned to the original experiment bottle. The number of uneaten copepods or *Artemia* in each feeding trial was counted. The feeding was calculated as the total number of [(copepods or Artemia) – (uneaten copepods or *Artemia*)].

We determined the growth rate during the exposure and recovery periods by measuring the body weight (g) of larva at each sampling point. Fish was weighted on a microbalance (accuracy of 0.1 mg, Satorious CPA224S, Germany).

### 2.5 Statistical analyses

Generalized mixed models (GMMs) were used to test single and interactive effects of Cu, MHW, food levels on survival, increased body weight, and feeding rate of *P. waigiensis* larvae during the exposure. We log-transformed the data log (x+1) of survival and feeding before analyses following the recommendation of Warton and Hui (2011). As the mortality was 100% in many treatments during the recovery period, we used simplified GMMs in which we included combinations of two stressors at a time: Cu and MHW, Cu and Food level, and MHW and Food level in the models. Data are presented in the graphs as means + 1 standard error (if not stated otherwise).

## 3. Results

### 3.1 Exposure period

Cu exposure strongly reduced the survival of fish larvae in a dose-dependent manner (main effects of Cu, Table 1, Figure 1). The lethal effect of Cu was particularly strong under MHW (Cu × MHW interaction, Table, Figure 1). None of the larvae survived at the highest Cu concentration (175 µg L^-1^) under MHW after 96h at low food and 144h at high food. MHW alone did not cause a lethal effect on fish larvae (Figure 1a,b).

**Table 1.**
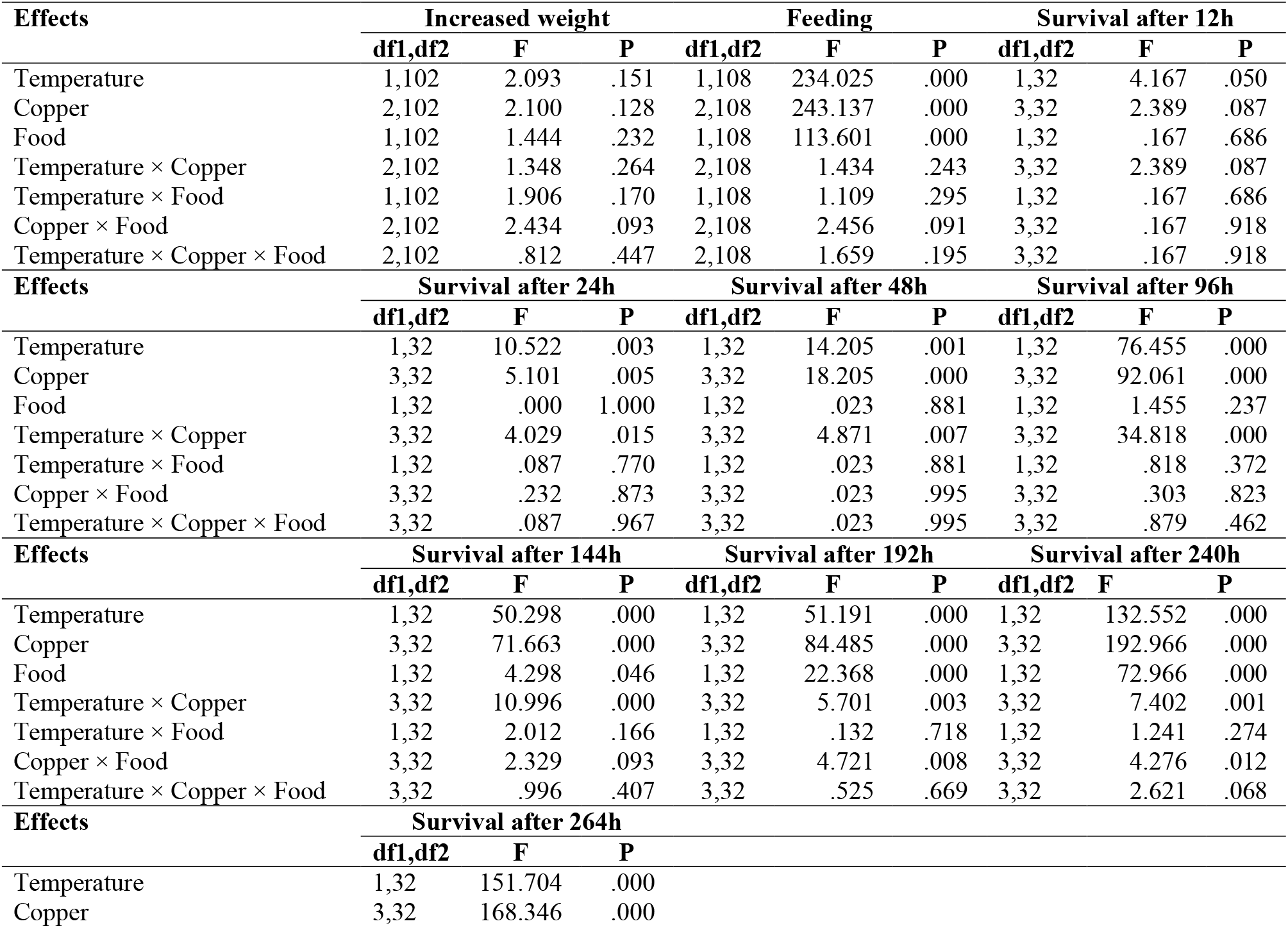
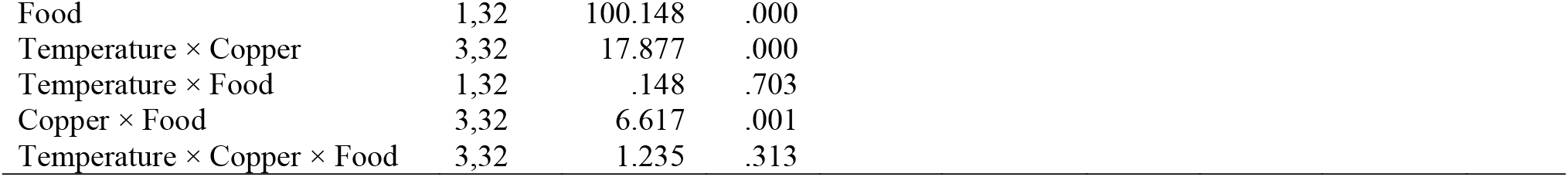
The statistical results of the generalized mixed models (GMMs) for the single and combined effects of Cu exposure, a simulated marine heatwave (MHW) and food limitation on survival, bodyweight and feeding of a reef-associated fish *Psammoperca waigiensis*.

| Effects | Increased weight |  |  | Feeding |  |  | Survival after 12h |  |  |
| --- | --- | --- | --- | --- | --- | --- | --- | --- | --- |
|  | df1,df2 | F | P | df1,df2 | F | P | df1,df2 | F | P |
| Temperature | 1,102 | 2.093 | .151 | 1,108 | 234.025 | .000 | 1,32 | 4.167 | .050 |
| Copper | 2,102 | 2.100 | .128 | 2,108 | 243.137 | .000 | 3,32 | 2.389 | .087 |
| Food | 1,102 | 1.444 | .232 | 1,108 | 113.601 | .000 | 1,32 | .167 | .686 |
| Temperature × Copper | 2,102 | 1.348 | .264 | 2,108 | 1.434 | .243 | 3,32 | 2.389 | .087 |
| Temperature × Food | 1,102 | 1.906 | .170 | 1,108 | 1.109 | .295 | 1,32 | .167 | .686 |
| Copper × Food | 2,102 | 2.434 | .093 | 2,108 | 2.456 | .091 | 3,32 | .167 | .918 |
| Temperature × Copper × Food | 2,102 | .812 | .447 | 2,108 | 1.659 | .195 | 3,32 | .167 | .918 |
| Effects | Survival after 24h |  |  | Survival after 48h |  |  | Survival after 96h |  |  |
|  | df1,df2 | F | P | df1,df2 | F | P | df1,df2 | F | P |
| Temperature | 1,32 | 10.522 | .003 | 1,32 | 14.205 | .001 | 1,32 | 76.455 | .000 |
| Copper | 3,32 | 5.101 | .005 | 3,32 | 18.205 | .000 | 3,32 | 92.061 | .000 |
| Food | 1,32 | .000 | 1.000 | 1,32 | .023 | .881 | 1,32 | 1.455 | .237 |
| Temperature × Copper | 3,32 | 4.029 | .015 | 3,32 | 4.871 | .007 | 3,32 | 34.818 | .000 |
| Temperature × Food | 1,32 | .087 | .770 | 1,32 | .023 | .881 | 1,32 | .818 | .372 |
| Copper × Food | 3,32 | .232 | .873 | 3,32 | .023 | .995 | 3,32 | .303 | .823 |
| Temperature × Copper × Food | 3,32 | .087 | .967 | 3,32 | .023 | .995 | 3,32 | .879 | .462 |
| Effects | Survival after 144h |  |  | Survival after 192h |  |  | Survival after 240h |  |  |
|  | df1,df2 | F | P | df1,df2 | F | P | df1,df2 | F | P |
| Temperature | 1,32 | 50.298 | .000 | 1,32 | 51.191 | .000 | 1,32 | 132.552 | .000 |
| Copper | 3,32 | 71.663 | .000 | 3,32 | 84.485 | .000 | 3,32 | 192.966 | .000 |
| Food | 1,32 | 4.298 | .046 | 1,32 | 22.368 | .000 | 1,32 | 72.966 | .000 |
| Temperature × Copper | 3,32 | 10.996 | .000 | 3,32 | 5.701 | .003 | 3,32 | 7.402 | .001 |
| Temperature × Food | 1,32 | 2.012 | .166 | 1,32 | .132 | .718 | 1,32 | 1.241 | .274 |
| Copper × Food | 3,32 | 2.329 | .093 | 3,32 | 4.721 | .008 | 3,32 | 4.276 | .012 |
| Temperature × Copper × Food | 3,32 | .996 | .407 | 3,32 | .525 | .669 | 3,32 | 2.621 | .068 |
| Effects | Survival after 264h |  |  |  |  |  |  |  |  |
|  | df1,df2 | F | P |  |  |  |  |  |  |
| Temperature | 1,32 | 151.704 | .000 |  |  |  |  |  |  |
| Copper | 3,32 | 168.346 | .000 |  |  |  |  |  |  |

|  |  |  |  |
| --- | --- | --- | --- |
| Food | 1,32 | 100.148 | .000 |
| Temperature × Copper | 3,32 | 17.877 | .000 |
| Temperature × Food | 1,32 | .148 | .703 |
| Copper × Food | 3,32 | 6.617 | .001 |
| Temperature × Copper × Food | 3,32 | 1.235 | .313 |

**Figure 1.**
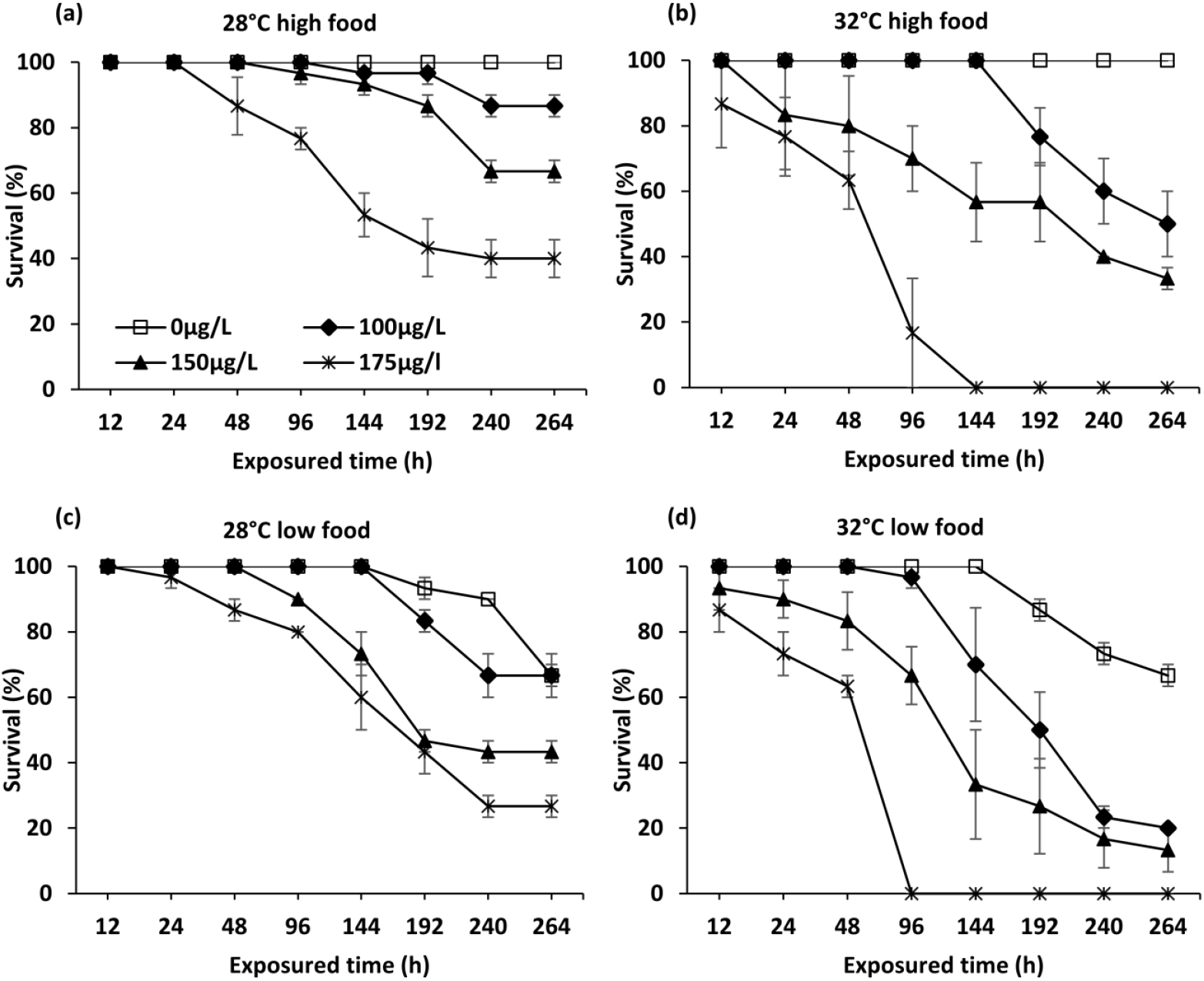
Survival of Waigieu seaperch (*Psammoperca waigiensis*) in response to Cu, marine heatwave and food limitation.

In the absence of Cu, food limitation directly resulted in ∼33% mortality of fish larvae, regardless of temperatures (Figure 1). Food limitation further exaggerated the effects of Cu, particularly under MHW (Cu × MHW × Food interaction, Table 1, Figure 1).

Under the high food level, Cu and MHW exposure reduced 3-5 times larval growth rate; the effect of Cu was stronger under MHW (Cu, MHW effects, Cu × MHW and Cu × MHW × Food interactions, Table 1, Figure 2A). The larval growth rate was extremely slow under food limitation (main effect of Food, Table 1), and there were unclear, inconsistent trends of the MHW interacting with food limitation and Cu (Figure 2B).

**Figure 2.**
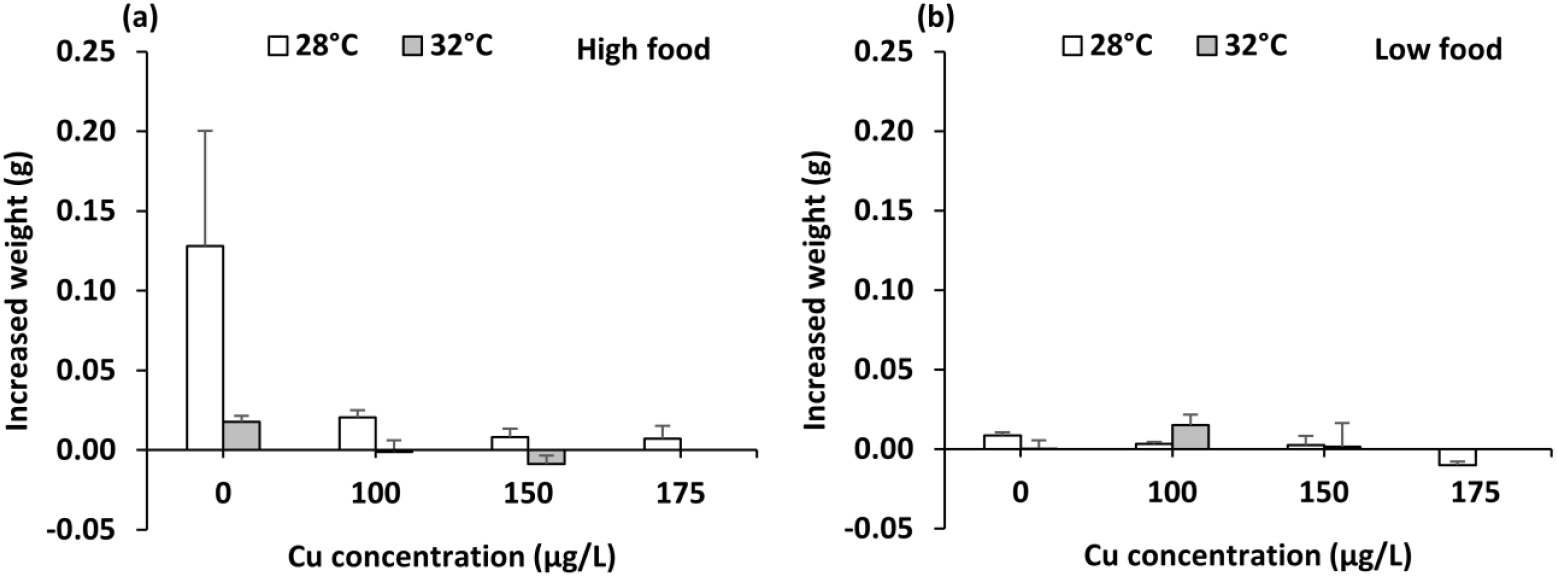
Changes in the bodyweight of Waigieu seaperch (*Psammoperca waigiensis*) in response to Cu, marine heatwave, and food limitation.

The feeding rate of larvae was slower in Cu, MHW and low food treatment (main effects of Cu, MHW and Food, Table 1, Figure 3). There were no two- and three-way interactions of Cu, MHW, and food treatment on the feeding rate of larvae (Table 1, Figure 3).

**Figure 3.**
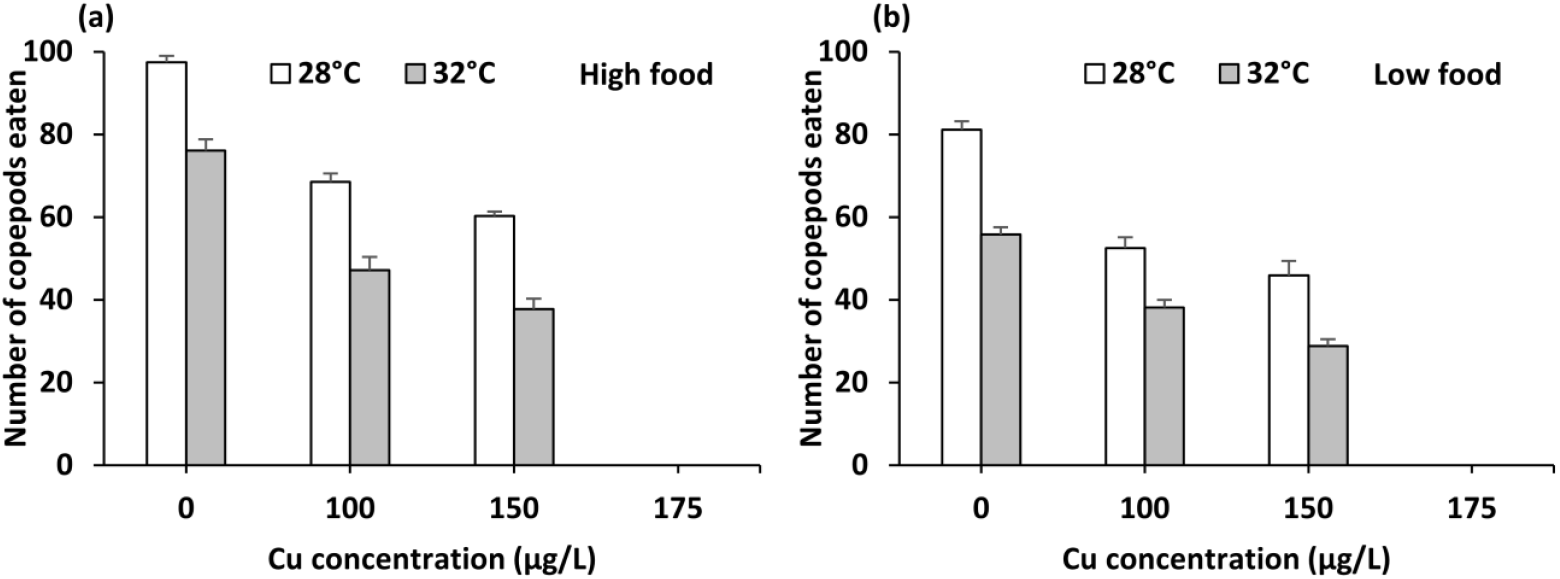
The feeding of Waigieu seaperch (*Psammoperca waigiensis*) in response to Cu, marine heatwave, and food limitation.

### 3.2 Post-exposure period

Survival was 100% in the control treatment (28°C, 0 µg Cu L^-1^, and high food). Mortality occurred in all other treatments, quicker and more pronounced in previously Cu-exposed larvae under MHW, and limited food (Table 2, Figure 5). In high food and 100 µg Cu L^-1^, survival was 9 times higher in well-fed larvae than in low-fed larvae (Table 2, Figure 5). None of the larvae survived in high food at 150 and 175 µg Cu L^-1^ and all previously exposed-Cu treatments under food limitation (Figure 4).

**Table 2.**
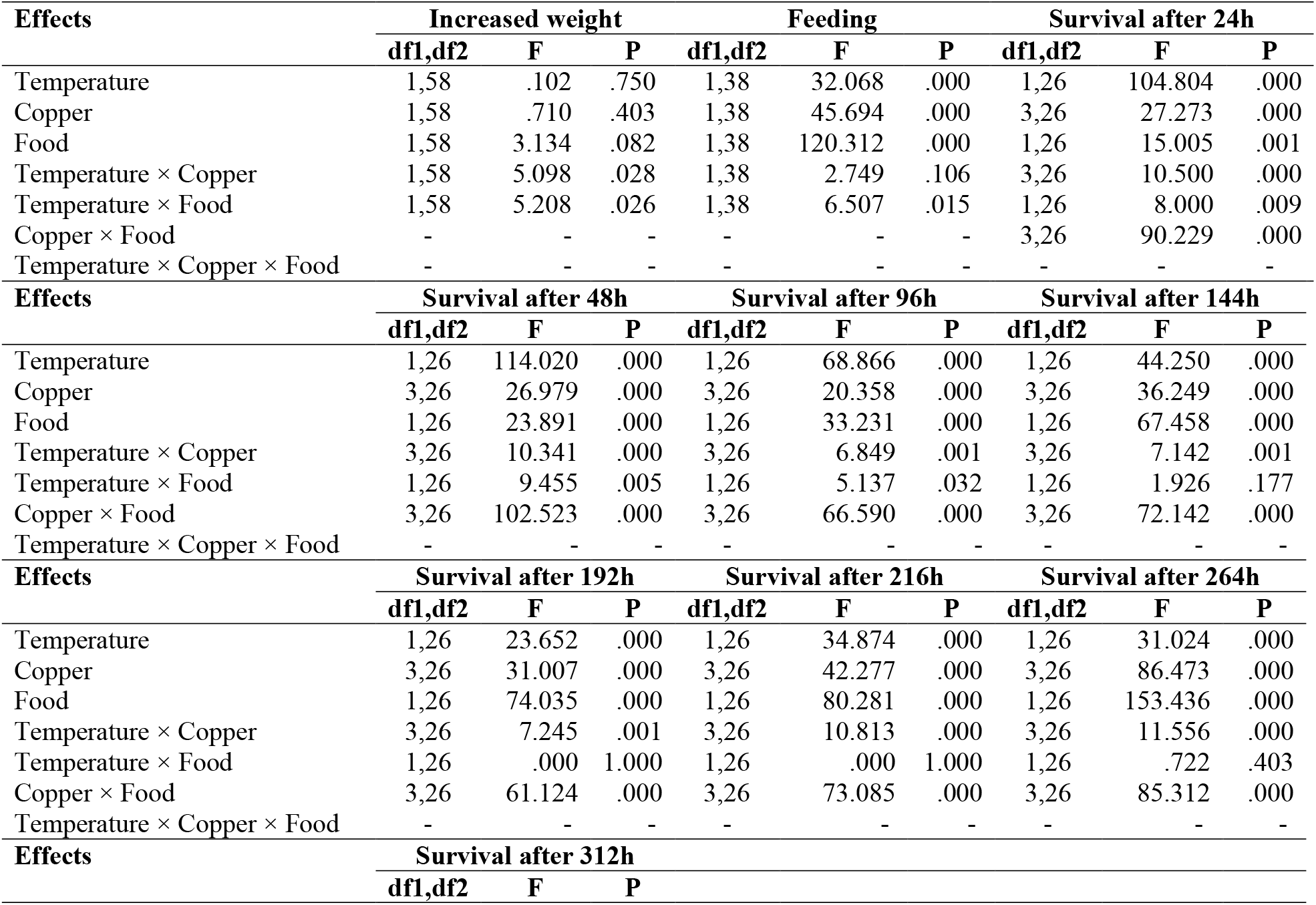
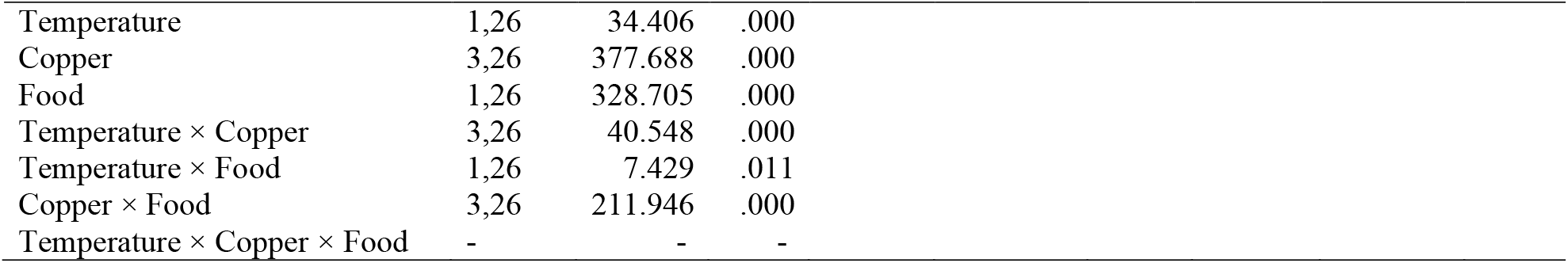
The statistical results of the generalized mixed models (GMMs) for the single and combined delayed effects of Cu exposure, a simulated marine heatwave (MHW) and food limitation on survival, body weight and feeding of a reef-associated fish *Psammoperca waigiensis*.

| Effects | Increased weight |  |  | Feeding |  |  | Survival after 24h |  |  |
| --- | --- | --- | --- | --- | --- | --- | --- | --- | --- |
|  | df1,df2 | F | P | df1,df2 | F | P | df1,df2 | F | P |
| Temperature | 1,58 | .102 | .750 | 1,38 | 32.068 | .000 | 1,26 | 104.804 | .000 |
| Copper | 1,58 | .710 | .403 | 1,38 | 45.694 | .000 | 3,26 | 27.273 | .000 |
| Food | 1,58 | 3.134 | .082 | 1,38 | 120.312 | .000 | 1,26 | 15.005 | .001 |
| Temperature × Copper | 1,58 | 5.098 | .028 | 1,38 | 2.749 | .106 | 3,26 | 10.500 | .000 |
| Temperature × Food | 1,58 | 5.208 | .026 | 1,38 | 6.507 | .015 | 1,26 | 8.000 | .009 |
| Copper × Food | - | - | - | - | - | - | 3,26 | 90.229 | .000 |
| Temperature × Copper × Food | - | - | - | - | - | - | - | - | - |
| Effects | Survival after 48h |  |  | Survival after 96h |  |  | Survival after 144h |  |  |
|  | df1,df2 | F | P | df1,df2 | F | P | df1,df2 | F | P |
| Temperature | 1,26 | 114.020 | .000 | 1,26 | 68.866 | .000 | 1,26 | 44.250 | .000 |
| Copper | 3,26 | 26.979 | .000 | 3,26 | 20.358 | .000 | 3,26 | 36.249 | .000 |
| Food | 1,26 | 23.891 | .000 | 1,26 | 33.231 | .000 | 1,26 | 67.458 | .000 |
| Temperature × Copper | 3,26 | 10.341 | .000 | 3,26 | 6.849 | .001 | 3,26 | 7.142 | .001 |
| Temperature × Food | 1,26 | 9.455 | .005 | 1,26 | 5.137 | .032 | 1,26 | 1.926 | .177 |
| Copper × Food | 3,26 | 102.523 | .000 | 3,26 | 66.590 | .000 | 3,26 | 72.142 | .000 |
| Temperature × Copper × Food | - | - | - | - | - | - | - | - | - |
| Effects | Survival after 192h |  |  | Survival after 216h |  |  | Survival after 264h |  |  |
|  | df1,df2 | F | P | df1,df2 | F | P | df1,df2 | F | P |
| Temperature | 1,26 | 23.652 | .000 | 1,26 | 34.874 | .000 | 1,26 | 31.024 | .000 |
| Copper | 3,26 | 31.007 | .000 | 3,26 | 42.277 | .000 | 3,26 | 86.473 | .000 |
| Food | 1,26 | 74.035 | .000 | 1,26 | 80.281 | .000 | 1,26 | 153.436 | .000 |
| Temperature × Copper | 3,26 | 7.245 | .001 | 3,26 | 10.813 | .000 | 3,26 | 11.556 | .000 |
| Temperature × Food | 1,26 | .000 | 1.000 | 1,26 | .000 | 1.000 | 1,26 | .722 | .403 |
| Copper × Food | 3,26 | 61.124 | .000 | 3,26 | 73.085 | .000 | 3,26 | 85.312 | .000 |
| Temperature × Copper × Food | - | - | - | - | - | - | - | - | - |
| Effects | Survival after 312h |  |  |  |  |  |  |  |  |
|  | df1,df2 | F | P |  |  |  |  |  |  |

|  |  |  |  |
| --- | --- | --- | --- |
| Temperature | 1,26 | 34.406 | .000 |
| Copper | 3,26 | 377.688 | .000 |
| Food | 1,26 | 328.705 | .000 |
| Temperature × Copper | 3,26 | 40.548 | .000 |
| Temperature × Food | 1,26 | 7.429 | .011 |
| Copper × Food | 3,26 | 211.946 | .000 |
| Temperature × Copper × Food | - | - | - |

**Figure 4.**
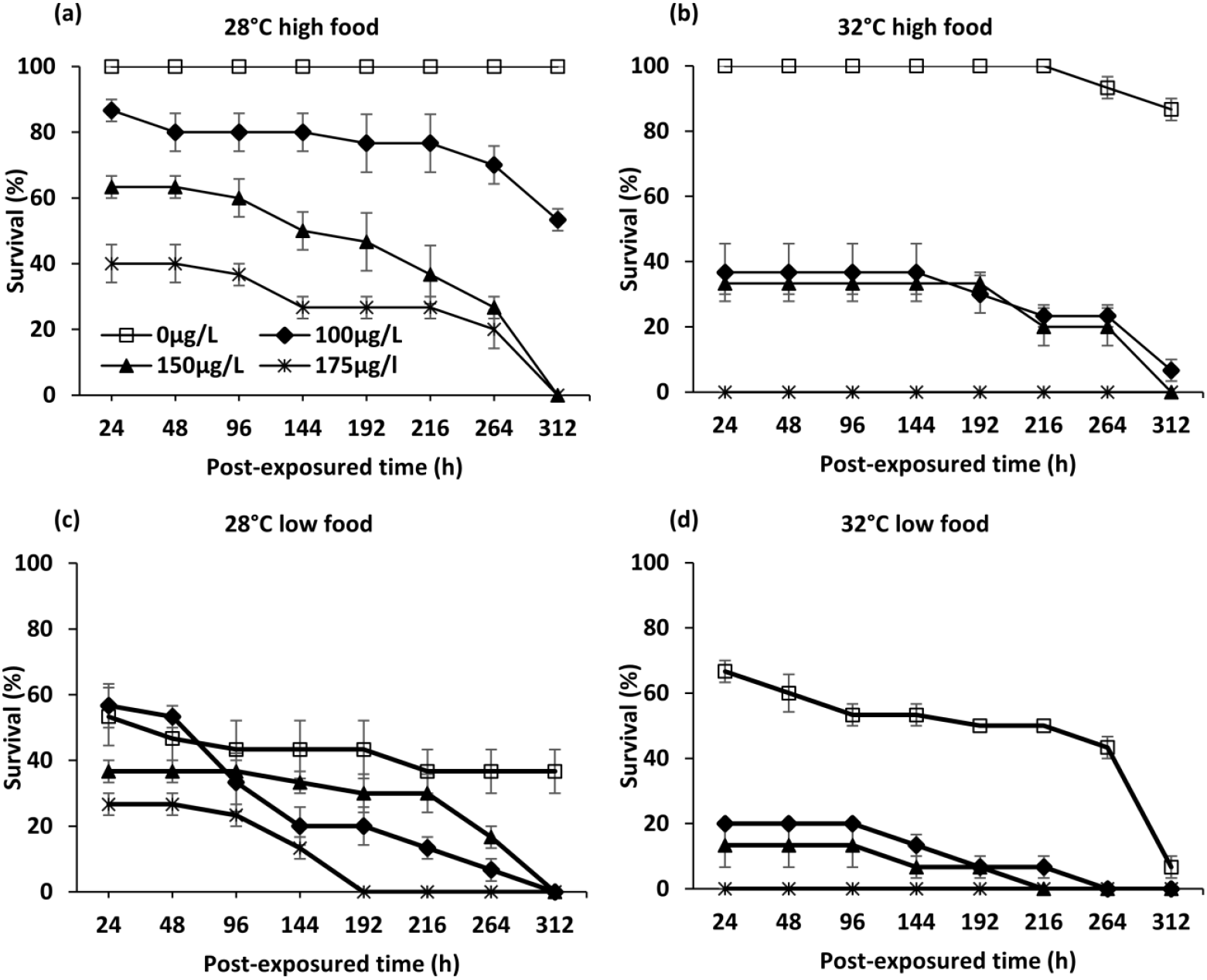
Delayed interactive effects of Cu exposure, marine heatwave, and food limitation on the survival of Waigieu seaperch (*Psammoperca waigiensis*) post-exposure period.

**Figure 5.**
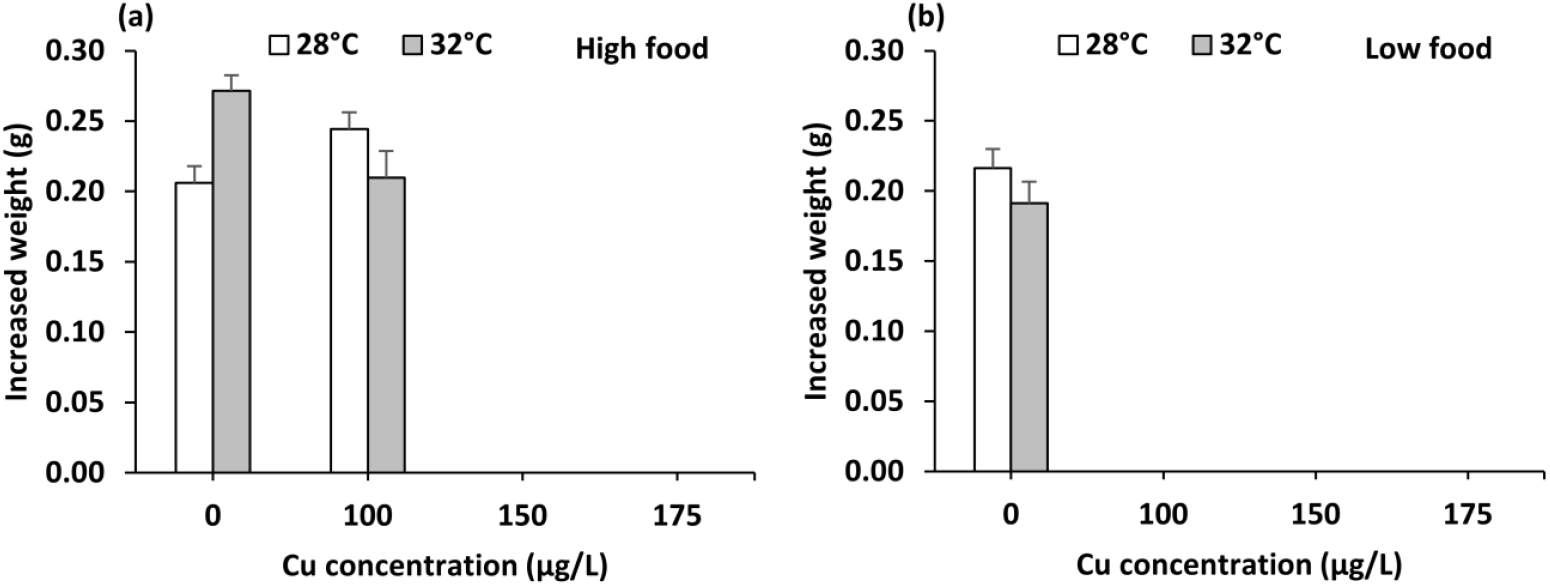
Delayed effects of Cu, marine heatwave and food limitation on the bodyweight of *Psammoperca waigiensis* larvae.

Overall, the bodyweight of fish larvae, in all treatments, increased several times higher than during the exposure period. In high food treatment, larvae showed the highest increased body weight in those which were previously exposed to MHW, followed by those who were previously exposed to 100 µg Cu L^-1^ at 28°C, while there was no difference in the control and larvae which were previously exposed to both 100 µg Cu L^-1^ and MHW (Table2, Figure 5a). In the low food treatment, previously MHW-exposed fish had a lower growth rate than those under control temperature (Table 2, Figure 5b).

The feeding was still substantially lower in previously Cu- and MHW-exposed larvae than those in control, especially in those who were previously exposed to both Cu and MHW simultaneously (Figure 6a). In low food treatment, the feeding rate of fish was ca. 40% of the control treatment (28°C, no Cu and high food). Yet, previously MHW-exposed larvae had 6 times lower feeding than those in the control treatment (Table 2, Figure 6b).

**Figure 6.**
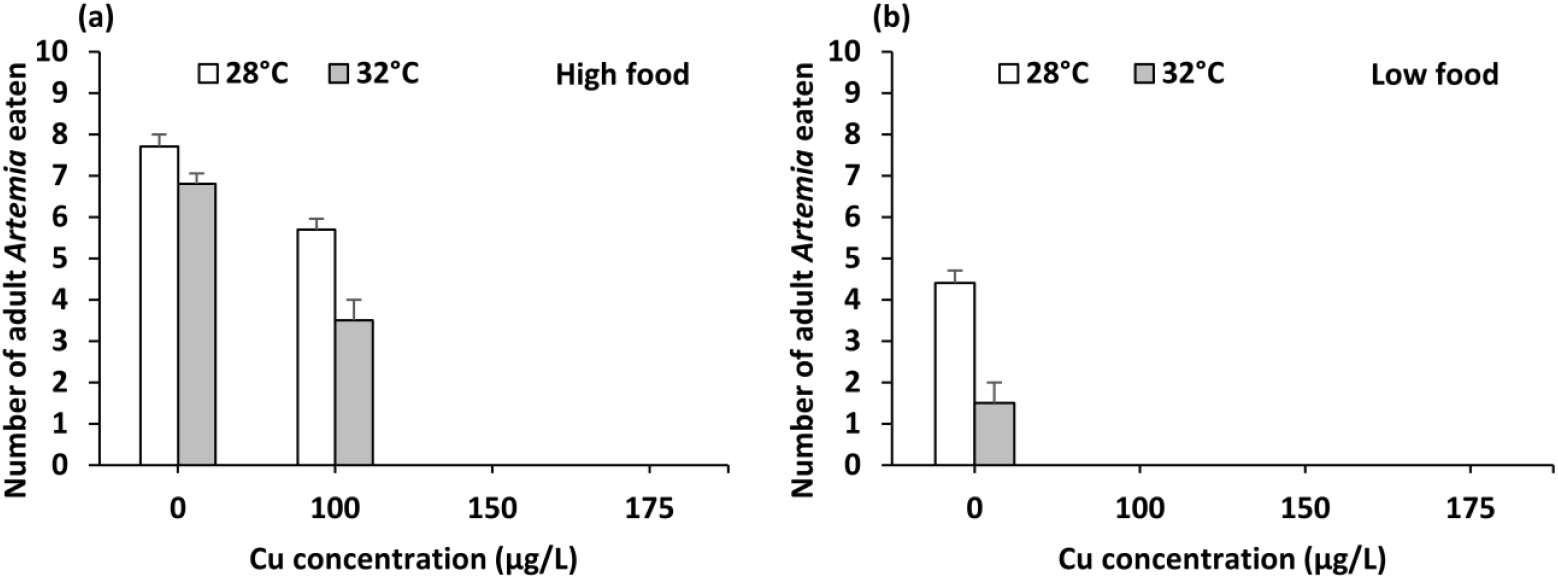
Delayed effects of Cu, marine heatwave and food limitation on the feeding of *Psammoperca waigiensis* larvae.

## 4. Discussion

### 4.1 Direct effects of Cu, marine heatwaves, food limitation and their interactions

We found strong and clear dose-dependent effects of Cu on all measured response variables of *P. waigiensis* larvae, including reduced survival, growth rate, and feeding. Cu is highly toxic to fish as it can increase cellular oxidative damages while impairing enzyme activities and repairing process and disrupting homeostasis (Baldissera et al., 2020; Braz-Mota et al., 2017; Xu et al., 2018), impairing behaviours (McIntyre et al., 2012; Sovova et al., 2014), and increased mortality (Braz-Mota et al., 2017). This is alarming as in this study, the exposure Cu concentrations of 100 - 175 µg Cu L^-1^ are lower than a limited Cu concentration of 200 µg L^-1^ allowed for the surface water for aquaculture purposes in Vietnam (MONRE-Vietnam, 2015) and fall within the measured Cu concentrations in coastal waters in the tropics (Jonathan et al., 2011; Le and Nguyen, 2018).

Importantly, MHW substantially increased the lethal effect of Cu on *P. waigiensis* larvae. All MHW-exposed larvae died at the highest Cu concentration, regardless of the food status. This is in agreement with the generally higher toxicity of metals with increasing temperature (Braz-Mota et al., 2017; Sokolova and Lannig, 2008) caused by impairing energy metabolism and increased oxidative stress (Sokolova and Lannig, 2008; Braz-Mota et al., 2017; Dinh Van et al., 2013; Fonseca et al., 2017). The higher toxicity of Cu under warming has also been observed before in marine zooplankton such as copepods collected at the same study location (Dinh et al., 2020b) or other fish species (Braz-Mota et al., 2017; Zebral et al., 2019), which may be an important factor to consider for ecological risk assessment as a scientific base for national and regional regulatory updates.

We did not find the direct lethal effect of MHW on *P. waigiensis* larvae. The lethal effect of MHWs on tropical fish larvae has been observed before, e.g. cobia *Rachycentron canadum* (Le et al., 2020). It seems that *P. waigiensis* larvae could cope better with heat stress than cobia larvae, which may be due to their smaller body size. The thermal tolerance of aquatic animals is generally determined by the capacity for oxygen supply in relation to oxygen demand (Pörtner, 2010; Pörtner et al., 2017; Verberk et al., 2016), which is disproportionate with the larger size of marine fish (Messmer et al., 2017). MHW substantially reduced the growth rate of fish larvae, which three non-exclusive mechanisms can explain: lower feeding, lower digestibility and higher energy expenditure to deal with heat stress. Indeed, larval feeding was lower under MHW and this pattern was consistent across Cu concentrations and food levels. This lowered feeding may be further enhanced by the lower protein and lipid digestibility. Indeed, our recent study has shown that MHW reduces the lower protein and lipid conversion efficiency of *P. waigiensis* larvae, thereby lowering energy intake, energy reserve and building block materials for the individual growth such as amino acids and proteins (Le et al., 2021). Finally, energy demand and expenditure for dealing with heat stress may be higher under MHW e.g., the upregulation of heat shock proteins (Li et al., 2015; Sørensen et al., 2003; Veilleux et al., 2018).

Food limitation caused the death of one-third population and reduced the growth rate (indicated by the bodyweight) from 3 - 5 times. Tropical fish larvae typically have a high metabolism and low energy reserve (Clarke and Johnston, 1999; Johnston and Battram, 1993) as the response to accelerate their early growth (Henderson, 2005; Metcalfe and Monaghan, 2001) which is critical for early survival from competition (Nakayama and Fuiman, 2010) and antipredator (Hoey and McCormick, 2004; Ziegelbecker and Sefc, 2021). It is likely that high mortality of fish larvae in low-food treatment may reflect their poor energy reserves such as lower lipid and protein contents (see e.g., in rainbow trout Josephson et al., 2012), which could not buffer them during a high energy demand period of a MHW to cope with heat stress. Across all Cu and MHW conditions, surviving larvae from food limitation showed a substantially reduced growth rate, 3-5 times compared to the control one. A reduced growth rate is a universe response of organisms to food shortage, resulting from a lowered energy intake while they may still have to spend energy expenditure on vital physiological and behavioural functions.

### 4.2 Post-exposure period

The delayed effects of Cu, MHW and food limitation on survival were manifested (for MHW), prolonged and/or magnified during the post-exposure periods. The delayed effect of Cu killed all remaining fish larvae at 150 and 175 µg L^-1^, and all larvae at 100 µg L^-1^ in low food. The strong delayed synergistic effect of Cu was present as indicated by a 9-time higher larval mortality of those previously exposed to MHW, particularly in food limitation than in the control treatment. Only one previous study has found a strong delayed synergistic effect of starvation and heatwaves together and contaminant (pesticide chlopyrifos) on an aquatic insect (*Coenagrion puella*), which was explained by the metabolic depression (Dinh et al., 2016a). A similar mechanism has not been explored for coastal marine fish, but metabolism and the capacity for oxygen supply in relation to oxygen demand are key factors determining the tolerance of fish to stressors such as warming (Pörtner, 2010; Pörtner et al., 2017; Pörtner and Farrell, 2008). Note that, we could not perform physiological analyses e.g., energy reserves, levels of heat shock proteins, oxidative stress and electron transport system as an indication for the cellular metabolism and oxygen consumption (see e.g., Dinh et al., 2016a) due to the low survival in many treatments, precluding the meaningful comparisons.

Surprisingly, MHW alone was not lethal during the exposure period, but it resulted in moderate mortality (approximately 15%) of fish larvae during the post-exposure period. The fastest acceleration of the compensation growth of previously HMH-exposed larvae may be an immediate cost of this mortality (see further below in the growth rate). The physiological mechanisms such as the overcompensation for the high metabolism to accelerate the growth which may be outpace the oxygen delivery (Portner et al., 2017; Pörtner, 2010; Pörtner and Farrell, 2008). The delayed effect of the heatwave on cellular metabolism and oxygen consumption has been observed before in aquatic insects (Dinh et al., 2016a). Irrespective of the physiological mechanisms for the delayed synergistic effects of Cu, MHW and starvations, strong delayed mortality of a reef-associate fish like *P. waigiensis* highlights the complexity of the stressor interactions (Jackson et al., 2021), which may accelerate the biodiversity loss in the tropical ecosystems (Barlow et al., 2018; Darling and Côté, 2008; Hughes et al., 2019).

Growth rate was rapidly recovered in all surviving larvae. The growth rate was fastest in those that were previously exposed to MHW under high food level without Cu while there were not much differences in other treatments. The recovery in growth rate as indicated by the increased larval weight, which can be explained by the compensation growth (reviewed in Dmitriew, 2011), even faster than the control group (see e.g., in Atlantic salmon juveniles Nicieza and Metcalfe, 1997). The compensation growth has been observed widespread across taxa (Dmitriew, 2011; Nicieza and Metcalfe, 1997). The increased food intake has been explained as the major mechanism for the compensation growth (see e.g., Calow, 1977; Dmitriew, 2011; Nicieza and Metcalfe, 1997), but the feeding of *P. waigiensis* was still lower in those which were previously exposed to MHW or Cu, particularly in previously-starved larvae. It suggests that a prioritize of energy acquisition on growth rate may have occurred, which may have various immediate and long-term costs (Dmitriew, 2011). Indeed, mortality occurred post-stress period was an example of the direct cost.

### 4.3 Applications and perspectives

The complex interactions of dominant stressors in the coastal marine ecosystems such as pollution, MHW and food shortage have been challenges when making projections of sea and ocean life in response to the present-day and future conditions. Our study contributes to tackling these challenges by revealing three important patterns crucial for ecological risk assessments (ERAs) of multiple stressors such as metals and MHW on the early life of tropical coastal fish. Firstly, we observed a strong synergistic effect of Cu and MHW, dominant stressors in the coastal environment on survival and growth rate, key parameters for the persistence of fish larvae. Secondly, we highlighted the role of food availability in modulating the interactive effects. Thirdly, the delayed synergistic effects of contaminants, MHW and starvation lasted beyond the exposure period, which are critically important to understand how ecological memory (see Hughes et al., 2019; Jackson et al., 2021) can accelerate biodiversity loss in the tropics (Barlow et al., 2018; Darling and Côté, 2008; Hughes et al., 2019). Together with our previous studies on zooplankton (Dinh et al., 2020a; Dinh et al., 2021; Dinh et al., 2020b), the lethal effects of Cu on fish larvae occur at a concentration that is lower than the Cu concentration allowed for the coastal water of Vietnam, 200 µg Cu/L (QCVN-10-MT-2015-BTNMT, MONRE-Vietnam, 2015), which urgently asks for the updates in local and national regulations to protect the marine, particularly reef environment. This is especially important and urgent as reefs and, more generally, coastal marine ecosystems in the Southeast Asian countries are experiencing increasing levels of various anthropogenic pollutants (Dinh, 2019; Jambeck et al., 2015; Nguyen et al., 2020) and marine heatwaves (Yao et al., 2020).

## Author contributions

M.H.L, K.V.D., X.T.V., and Q.H.P designed the experiment; X.T.V. and M.H.L. conducted the experiment, collected and analysed the data. M.H.L and K.V.D wrote the first draft of the manuscript. All authors contributed to the later version of the manuscript; all authors read and approved the manuscript for publication.

## Acknowledgements

This research is funded by Vietnam National Foundation for Science and Technology Development (NAFOSTED) under grant number 106.05-2017.343.

